# Stage-dependent chromatin accessibility remodeling defines an architectural state in facioscapulohumeral muscular dystrophy myoblasts

**DOI:** 10.64898/2026.03.06.710110

**Authors:** Francesca Losi, Bruno Fosso, Rossella Tupler, Valentina Salsi

## Abstract

Facioscapulohumeral muscular dystrophy (FSHD) has been linked to alterations in higher-order genome organization, yet how these structural perturbations shape chromatin accessibility across muscle developmental stages remains unclear. Here, we identify a stage-specific architectural chromatin accessibility state that defines proliferating FSHD myoblasts. Using genome-wide ATAC-seq profiling of primary human myoblasts and differentiated myotubes, analyzed using the telomere-to-telomere (T2T-CHM13) human reference genome, we map the genome-wide distribution and hierarchical organization of this state.

We detect extensive chromatin accessibility remodeling, with over 12,000 differentially accessible regions in myoblasts and marked attenuation of this state upon terminal differentiation. Remodeling predominantly affects intronic and distal intergenic regions enriched in non-coding and repeat-proximal sequences, indicating redistribution of regulatory accessibility beyond promoter-centered regulation.

At the chromosome scale, accessibility losses cluster within megabase domains enriched in heterochromatin- and nucleolus-associated regions, revealing coordinated reorganization of nuclear architecture. No reproducible accessibility changes occur within the D4Z4 repeat array or canonical DUX4 target loci, demonstrating that early architectural remodeling in FSHD myoblasts is largely uncoupled from sustained DUX4-driven programs.

These findings these findings define a transient architectural chromatin accessibility state that emerges in proliferating FSHD myoblasts and is largely resolved upon terminal differentiation.

Our work supports a model in which disruption of higher-order genome organization represents an early and stage-restricted determinant of disease susceptibility.

## Introduction

Facioscapulohumeral dystrophy (FSHD) is a hereditary muscle disorder characterized by progressive and variable muscle weakness, with marked inter-individual and intra-familial heterogeneity (Ricci et al. 2013; Ruggiero et al. 2020). While disease pathogenesis has traditionally been attributed to aberrant gene expression during muscle differentiation, accumulating evidence indicates that FSHD is fundamentally associated with disruption of higher-order genome organization and epigenetic regulation of repetitive DNA. In particular, contraction of the D4Z4 macrosatellite repeat at chromosome 4q35 leads to epigenetic alterations affecting nuclear architecture (Bodega et al. 2009), chromatin topology, (Gaillard et al. 2019; Robin et al. 2014) and repeat-associated regulatory networks (Salsi et al. 2025a; Cabianca et al. 2012).

Chromatin accessibility plays a central role in defining cellular competence and lineage-specific responses by constraining the regulatory landscape available to transcriptional programs. In skeletal muscle, dynamic chromatin remodeling accompanies the transition from proliferating myoblasts to differentiated myotubes, enabling the activation of myogenic genes while progressively restricting alternative transcriptional states (Asp et al. 2011; Hernández-Hernández et al. 2020). Perturbations of this process have been implicated in several neuromuscular disorders (Wilson 2000; Méjat and Misteli 2010; Dilworth and Blais 2011; Rugowska et al. 2021).

In FSHD epigenetic derepression of repetitive elements and alterations in nuclear genome organization (Gabellini 2004; Bodega et al. 2009; Robin et al. 2014; Gaillard et al. 2019; Salsi et al. 2020) suggests that disruption of chromatin architecture may occur early during muscle lineage progression (Ottaviani et al. 2009; Himeda et al. 2015). However, whether chromatin accessibility is altered at pre-differentiation stages in FSHD, prior to overt transcriptional dysregulation, remains unclear.

Consistent with this model, FSHD has long been recognized as a disorder with a strong epigenetic component (de Greef et al. 2009; Cabianca et al. 2012; Himeda et al. 2015; Nikolic et al. 2020). Chromosome conformation capture-based approaches and DNA-FISH analyses have shown that the 4q35 region engages in dynamic long-range interactions with other genomic loci and nuclear compartments (Bodega et al. 2009; Broucqsault et al. 2013; Robin et al. 2014; Gaillard et al. 2019; Laberthonnière et al. 2019) supporting the idea that disease-relevant alterations involve changes in nuclear organization rather than simple local regulatory defects. In parallel, transient expression of DUX family transcription factors has been shown to induce widespread chromatin accessibility changes through pioneer-like activity and recruitment of chromatin-modifying cofactors (Geng et al. 2012; Whiddon et al. 2017; Vuoristo et al. 2022). Yet, endogenous DUX4 expression is rare and mosaic in FSHD muscle cells, with DUX4-positive nuclei detected only in a small fraction of cells, including subsets of proliferating myoblasts and differentiated myotube nuclei (Snider et al. 2010; Jones et al. 2012; Tassin et al. 2013; Haynes et al. 2018). Consistent with this heterogeneity, single-cell and single-nucleus transcriptomic studies have identified distinct DUX4-high and DUX4-low nuclear populations, indicating that DUX4-driven transcriptional activation is not uniformly present across nuclei and may be variably detectable in bulk measurements (Jiang et al. 2020). Accordingly, comparative transcriptomic analyses have demonstrated that DUX4 target gene expression provides a variable and often weak molecular signature in patient samples, in contrast to more robust and reproducible biomarkers such as repression of PAX7 target genes (Banerji and Zammit 2019). Thus, these chromatin accessibility alterations can be temporally restricted and uncoupled from stable transcriptional output, indicating that chromatin architecture and regulatory competence may be altered independently of sustained transcriptional activation (Snider et al. 2010; Bosnakovski et al. 2014). These observations raise the possibility that pathogenic chromatin alterations in FSHD emerge as early architectural perturbations, rather than solely as secondary consequences of persistent DUX4-driven transcriptional programs.

To date, epigenomic investigations in FSHD have largely focused on transcriptional deregulation and epigenetic alterations associated with the D4Z4 locus and muscle differentiation, using both patient muscle biopsies and cultured muscle cells (Bodega et al. 2009; de Greef et al. 2009; Himeda et al. 2015; Jiang et al. 2020). However, the chromatin accessibility landscape of proliferating primary myoblasts, where early architectural and epigenetic alterations may arise prior to terminal differentiation, remains uncharacterized. Emerging evidence suggests that disease-relevant chromatin alterations may already be present in myoblasts (Stadler et al. 2013; Van Den Heuvel et al. 2019; Salsi et al. 2025b), potentially influencing cellular stress responses, lineage competence, and differentiation trajectories. Many disease-associated alterations are thought to arise from changes in chromatin repression and higher-order genome organization rather than from mutations affecting protein-coding genes (Masny et al. 2004; Bodega et al. 2009; Ottaviani et al. 2009; Salsi et al. 2025a). In this context, profiling chromatin accessibility across distinct stages of muscle differentiation may provide a powerful approach to distinguish early chromatin alterations from secondary transcriptional consequences.

The recent telomere-to-telomere (T2T) genome assemblies have further expanded the scope of epigenomic analyses by enabling systematic interrogation of repeat-rich, subtelomeric, and structurally complex genomic regions that are poorly represented in conventional genome builds (Miga and Sullivan 2021; Nurk et al. 2022; Haws et al. 2022; Salsi et al. 2026)

Here, we performed genome-wide ATAC-seq profiling (Buenrostro et al. 2015) in primary myoblasts and differentiated myotubes derived from FSHD patients and matched controls, using the T2T-CHM13 reference genome to enable systematic analysis of repeat-rich and structurally complex genomic regions. By directly comparing proliferative and differentiated states, we sought to determine whether FSHD is associated with stage-specific alterations in chromatin accessibility linked to higher-order genome organization, and to define when such architectural accessibility states emerge during muscle cell differentiation.

Our analyses reveal extensive chromatin accessibility remodeling in proliferating FSHD myoblasts, contrasted by a marked attenuation of differences in differentiated myotubes.

Notably, accessibility changes in myoblasts preferentially affect intergenic and heterochromatin-rich genomic regions rather than canonical muscle gene promoters, indicating that chromatin dysregulation in FSHD primarily reflects reorganization of regulatory and structural genome architecture. Together, these findings reveal the stage-specific chromatin accessibility landscape in FSHD and provide a framework for understanding how perturbations of nuclear organization precede and shape later transcriptional and cellular phenotypes.

## Results

### Stage dependent chromatin accessibility remodeling defines a distinct architectural state in FSHD myoblasts

To investigate chromatin accessibility alterations associated with FSHD across stages of muscle differentiation, we performed ATAC-seq in primary human myoblasts and corresponding differentiated myotubes derived from n = 2 FSHD patients and n = 2 unaffected controls (Supplementary_Figure_S1A). To minimize inter-individual variability and genetic background effects, patient and control samples were obtained from well-characterized familial cohorts, including affected and unaffected relatives, as previously described (Salsi et al. 2025a) and shown in Supplementary_Figure_S1B. This experimental design enabled identification of disease-associated chromatin accessibility differences within a shared genetic context. Multidimensional scaling analysis further supported the robustness of this experimental setting, showing clear separation between FSHD and control myoblasts, which served as the source population for the differentiated myotubes, while preserving inter-individual variability within the FSHD group (Supplementary Figure S1C).

Given the incomplete representation of structurally complex genomic regions in conventional human genome assemblies, all ATAC-seq analyses were performed using the telomere-to-telomere (T2T-CHM13v2.0) human reference genome. This approach enabled systematic assessment of chromatin accessibility across the full genomic landscape within a more complete and accurately annotated reference framework.

Chromatin accessibility was analyzed in proliferating myoblasts and terminally differentiated myotubes to distinguish stage-specific alterations from changes associated with myogenic maturation. Differentially accessible regions (DARs) were defined as genomic regions showing consistent accessibility differences between FSHD and control samples. To ensure robust identification of reproducible differences, DARs were identified by comparing all FSHD samples collectively to all controls and retaining regions exhibiting concordant directional changes across biological replicates. Statistical significance was determined using a genome-wide threshold of false discovery rate (FDR) < 0.05, following multiple testing correction, together with an absolute log_2_ fold change ≥ 1, ensuring identification of statistically significant and biologically meaningful accessibility differences. Regions consistently more accessible in FSHD were classified as Up-DARs, whereas regions showing reduced accessibility were classified as Down-DARs (Supplementary_Table_S1 and Figure 1A).

**Figure 1.**
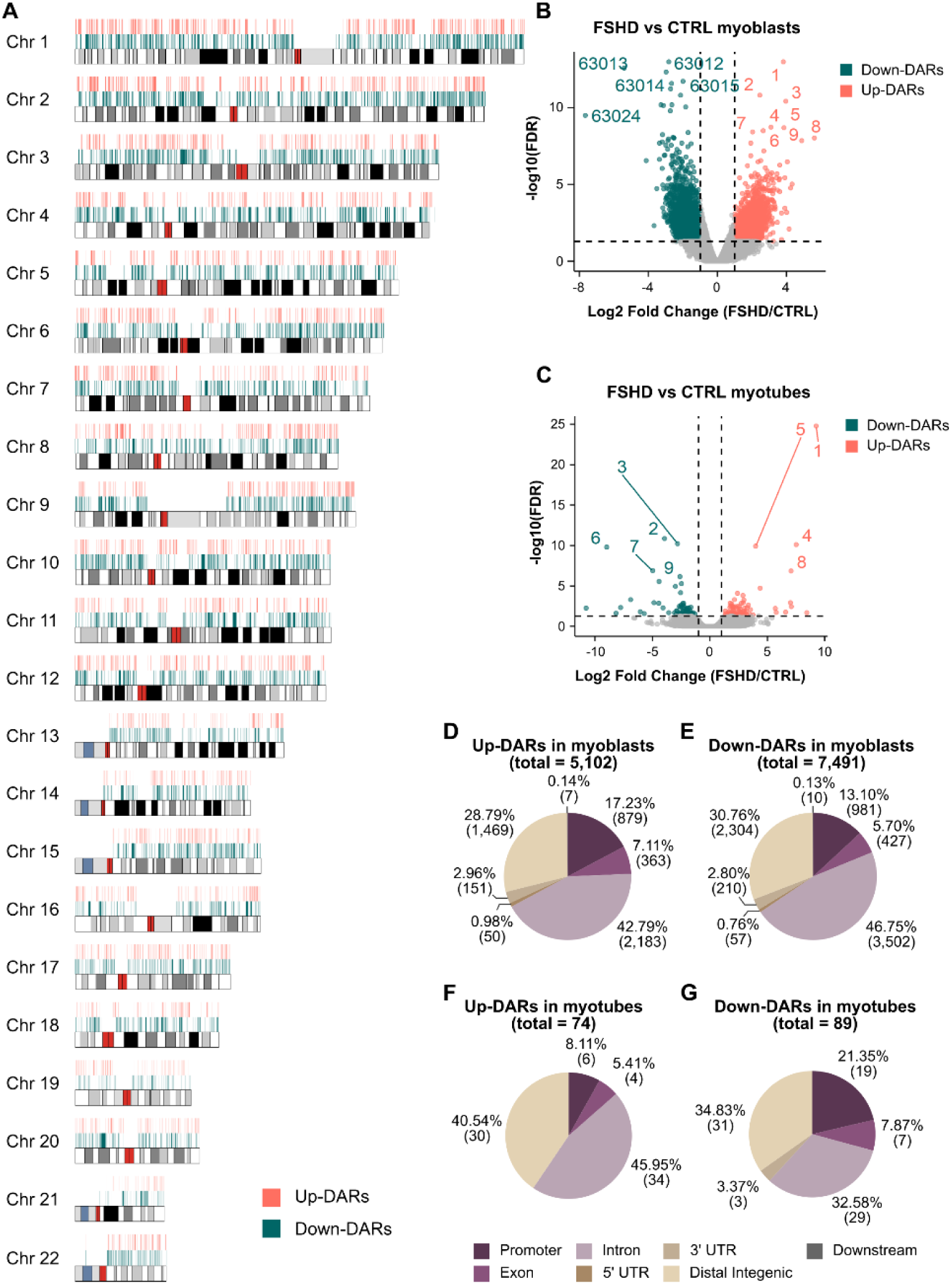
Genome-wide identification and genomic distribution of differentially accessible chromatin regions in FSHD myoblasts and myotubes. (A) Chromosome-wide distribution of differentially accessible regions across all human autosomes. Each horizontal track represents an individual chromosome, arranged from chromosome 1 to chromosome 22. Up-DARs are indicated in red and Down-DARs in teal. Black and gray segments represent cytogenetic bands (cytobands) along each chromosome. The positions of DARs are shown relative to chromosomal coordinates, illustrating their genome-wide distribution. (B) Volcano plot showing differential chromatin accessibility between FSHD and control primary human myoblasts. Each point represents an individual ATAC-seq peak. The x-axis shows the log₂ fold change in accessibility (FSHD/CTRL), and the y-axis shows the −log₁₀(FDR). Regions with significantly increased accessibility in FSHD (Up-DARs) are shown in red, and regions with significantly decreased accessibility (Down-DARs) are shown in teal. Vertical dashed lines indicate fold-change thresholds, and the horizontal dashed line indicates the statistical significance threshold. (***1***=chr19:42663366-42663803; ***2***=chr8:125236306-125237721; ***3***=chr4:37517813-37518483; ***4***=chr15:84909779-84910176; ***5***=chr7:146872319-146873047; ***6***=chr1:180272411-18027321; ***7***=chr7:41837735-41842398; ***8***=chr7:146578703-146579358; ***9***=chr7:147089937-147090658; ***63012***=chr7:37253326-37254866; ***63013***=chr8:11514184-11516273; ***63014***=chr5:55485814-55486163; ***63015***=chr21:30704676-30707177; ***63024***=chr8:11518982-11519321). (C) Volcano plot showing differential chromatin accessibility between FSHD and control myotubes, displayed as in panel B. Each point represents an individual ATAC-seq peak, with Up-DARs shown in red and Down-DARs shown in teal. (***1***=chr7:76053318-76053881; ***2***=chr1:16126339-16127460; ***3***=chr10:131229506-131231084; ***4***=chr22:21699861-21700117; ***5***=chr14:2770953-2771283; ***6***=chr9:80192322-80193046; ***7***=chr8:11514182-11516352; ***8*** = chr11:221023-221287; ***9***=chr3:69747056-69747802). (D) Genomic annotation of Up-DARs identified in human primary myoblasts (HPMs; total = 5,102). Peaks were classified based on genomic location relative to annotated gene features, including promoter, exon, intron, 5′ untranslated region (UTR), 3′ UTR, downstream region, and distal intergenic region. Percentages and absolute counts are indicated. (E) Genomic annotation of Down-DARs identified in HPMs (total = 7,491), classified according to genomic features as in panel C. (F) Genomic annotation of Up-DARs identified in myotubes (total = 74), categorized according to genomic features as described above. (G) Genomic annotation of Down-DARs identified in myotubes (total = 89), categorized according to genomic features as described above.

Using this approach, 12593 DARs were identified in proliferating myoblasts, of which 5102 showed increased accessibility (Up-DARs) and 7491 showed reduced accessibility (Down-DARs) (Figure 1B). In contrast, differentiated myotubes displayed fewer than 200 DARs in total, with substantially reduced effect sizes and no coherent global shifts in chromatin accessibility (Figure 1C). These results indicate that chromatin accessibility remodeling in FSHD is stage-dependent, with the vast majority of disease-associated changes occurring prior to terminal myogenic differentiation.

To define the genomic context of chromatin accessibility changes in FSHD myoblasts, DARs were annotated relative to gene features using the T2T-CHM13 reference genome. Across the genome, the majority of differential accessibility occurred outside canonical promoter regions. Specifically, 70-80% of both Up- and Down-DARs localized to intronic or distal intergenic regions, whereas only 13-17% overlapped annotated promoters, defined as -3 kb from the transcription start site (Figure 1D,E). These proportions were highly similar for regions showing increased or decreased accessibility, indicating that promoter-associated changes represent a minor fraction of the overall chromatin remodeling landscape.

In differentiated myotubes, the few regions retaining significant accessibility differences between FSHD and control samples (Figure 1F,G) did not define a coherent chromatin accessibility signature (Supplementary_Table_S1). These results define a stage-restricted accessibility signature in FSHD, with widespread remodeling in proliferating myoblasts and minimal residual differences in terminally differentiated myotubes.

### Chromatin accessibility remodeling preferentially targets non-coding regulatory space

To further define the genomic classes involved in chromatin accessibility remodeling in FSHD myoblasts, differentially accessible regions (DARs) were classified according to the biotype of their associated transcripts (Figure 2A,B). Across the dataset, non-coding loci accounted for a substantial fraction of differential accessibility, representing approximately 45–50% of all DARs. This proportion was comparable between regions showing increased and decreased accessibility, indicating that chromatin remodeling at non-coding loci is quantitatively extensive but not directionally biased toward either opening or closing.

**Figure 2.**
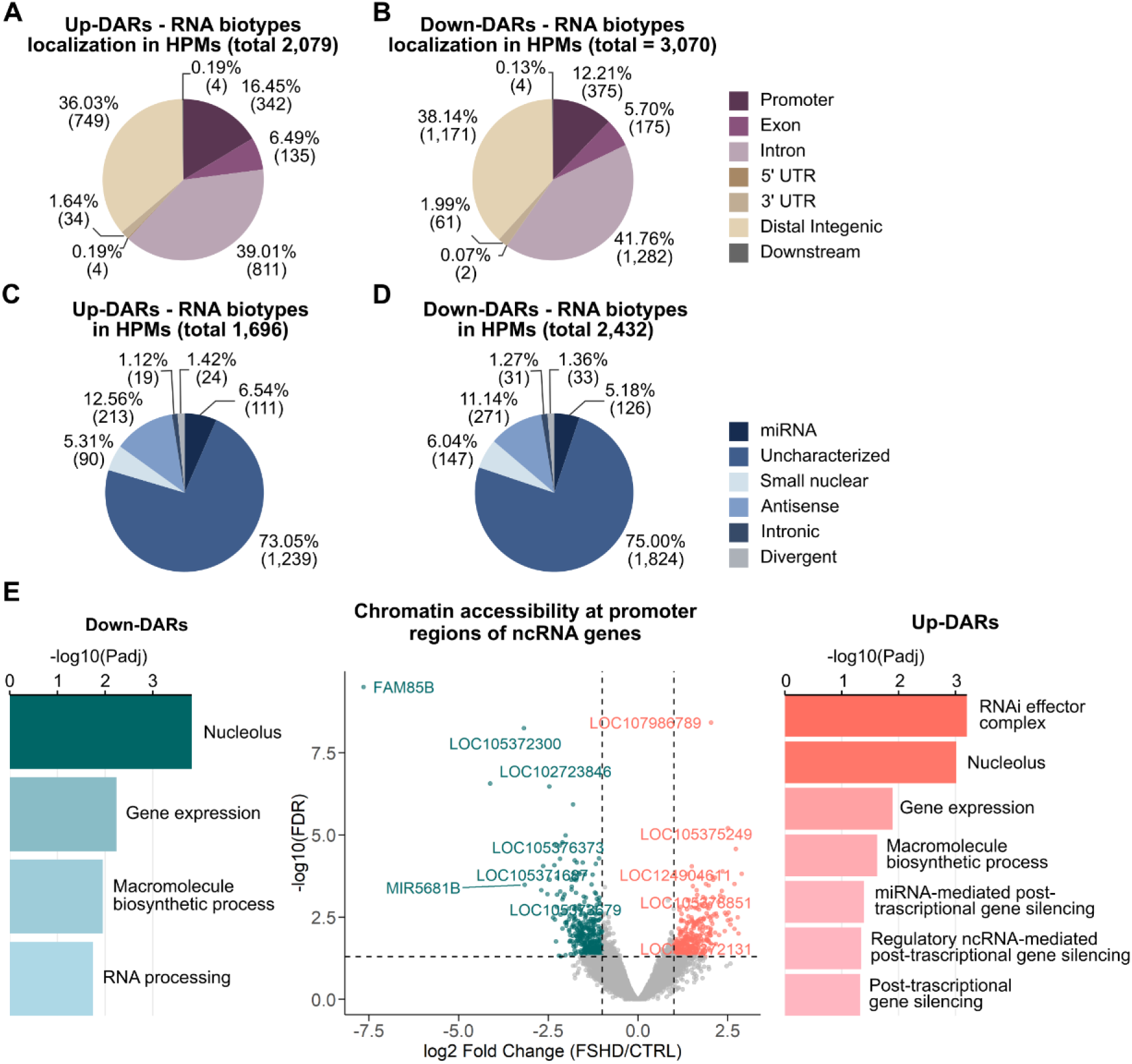
Genomic and functional annotation of differentially accessible regions associated with non-coding RNA loci in FSHD myoblasts. (A) Genomic distribution of Up-DARs relative to annotated non-coding gene features in human primary myoblasts (HPMs), restricted to regions overlapping RNA-associated loci (total = 2,079). Peaks were classified according to their genomic localization, including promoter, exon, intron, 5′ untranslated region (UTR), 3′ UTR, distal intergenic region, and downstream region. Percentages and absolute counts are indicated for each category. (B) Genomic distribution of Down-DARs relative to annotated gene features in HPMs (total = 3,070), categorized according to genomic location as described in panel A. Percentages and absolute counts are shown. (C) Distribution of Up-DARs overlapping annotated RNA biotypes in HPMs (total = 1,696). Peaks were assigned to RNA categories including miRNA, uncharacterized transcripts, small nuclear RNA, antisense RNA, intronic transcripts, and divergent transcripts. Percentages and absolute counts are indicated. (D) Distribution of Down-DARs overlapping annotated RNA biotypes in HPMs (total = 2,432), categorized according to RNA biotype as described in panel C. Percentages and absolute counts are shown. (E) Functional annotation of genes associated with promoter regions overlapping differentially accessible chromatin regions linked to non-coding RNA loci. Bar plots show selected Gene Ontology (GO) biological process terms associated with Down-DARs (left) and Up-DARs (right). The x-axis indicates the −log₁₀(adjusted P-value). The central panel shows a volcano plot of chromatin accessibility changes at promoter regions of ncRNA genes, with the x-axis representing log₂ fold change (FSHD/CTRL) and the y-axis representing −log₁₀(FDR). Each point represents an individual ncRNA-associated promoter region, with Up-DARs shown in red and Down-DARs shown in teal. Selected ncRNA loci are labeled.

Genomic annotation of non-coding DARs revealed that the large majority localized outside canonical promoter regions. For both Up- and Down-DARs, intronic and distal intergenic regions together accounted for approximately 75-80% of non-coding peaks, whereas only a minor fraction overlapped annotated non-coding promoters or exonic regions (Figure 2A,B). Enrichment analysis confirmed that DARs were significantly depleted from promoter regions and enriched in intronic and intergenic regions, indicating preferential chromatin accessibility remodeling outside canonical promoter regulatory regions (Table 1).

**Table 1.** Enrichment of differentially accessible regions across genomic features. Differentially accessible regions (DARs) were classified relative to promoter (-3 kb from transcription start site), intronic, and intergenic genomic regions using the T2T-CHM13 RefSeq annotation. Enrichment was assessed relative to the full ATAC-seq peak set using Fisher’s exact test. P-values were corrected for multiple testing using the Benjamini-Hochberg procedure. Odds ratios greater than 1 indicate enrichment, whereas odds ratios less than 1 indicate depletion. The “Result” column indicates enrichment status based on odds ratio and false discovery rate (FDR < 0.05).

| Set | Feature | Odds_Ratio | p_value | FDR | Result |
| --- | --- | --- | --- | --- | --- |
| **All_DARs** | Promoter | 0,676 | 4,13E-49 | 1,24E-48 | Depleted |
|  | Intron | 1,16 | 5,85E-16 | 8,78E-16 | Enriched |
|  | Intergenic | 1,07 | 1,50E-03 | 1,50E-03 | Enriched |
| **Up_DARs** | Promoter | 0,796 | 9,67E-09 | 2,90E-08 | Depleted |
|  | Intron | 1,14 | 7,53E-06 | 1,13E-05 | Enriched |
|  | Intergenic | 1,01 | 8,26E-01 | 8,26E-01 | Not changed |
| **Down_DARs** | Promoter | 0,597 | 1,35E-48 | 4,05E-48 | Depleted |
|  | Intron | 1,18 | 5,24E-12 | 7,86E-12 | Enriched |
|  | Intergenic | 1,11 | 7,51E-05 | 7,51E-05 | Enriched |

Stratification by non-coding RNA subtype revealed a strongly skewed composition. Intronic long non-coding RNAs represented approximately 75% of both Up- and Down-non-coding DARs, whereas antisense and divergent transcripts contributed smaller but consistent fractions. In contrast, small RNA classes, including microRNAs and snoRNAs, accounted for only a marginal proportion of non-coding DARs (5–7%)(Figure 2C,D). Notably, the relative contribution of each non-coding RNA subtype was nearly identical between accessibility gains and losses. Because a large fraction of non-coding DARs mapped to distal intergenic regions lacking clear gene annotations, functional interpretation of the full non-coding dataset using gene-centric enrichment approaches is inherently limited. We therefore restricted functional enrichment analysis to the subset of non-coding DARs overlapping annotated non-coding promoters (-3 kb from the transcription start site), representing genomic regions with a defined transcriptional anchor.

Although this promoter-associated subset constituted only a minority of non-coding DARs, enrichment analysis revealed reproducible associations with RNA-related functional categories. Both Up- and Down-regulated non-coding gene promoters were enriched for terms related to RNA processing, post-transcriptional gene silencing, RISC complex components, and nucleolar functions (adjusted p < 0.05; Figure 2E). These findings underscore the extensive involvement of a largely unexplored non-coding regulatory landscape in early-stage FSHD.

### Promoter accessibility remodeling reveals coordinated shifts in regulatory competence

Despite the predominance of chromatin accessibility changes at non-coding regulatory regions in FSHD myoblasts, we next examined promoter-associated accessibility changes at protein-coding genes to assess their contribution to regulatory competence and transcriptional reprogramming. Functional enrichment analysis of genes associated with differentially accessible promoter regions identified by ATAC-seq revealed distinct and directionally coherent regulatory signatures (Figure 3A). Promoter-associated peaks were stratified according to their distance from the transcription start site (TSS), with proximal promoters (≤1 kb) constituting the largest fraction of both Up- and Down-regulated groups (Figure 3B).

**Figure 3.**
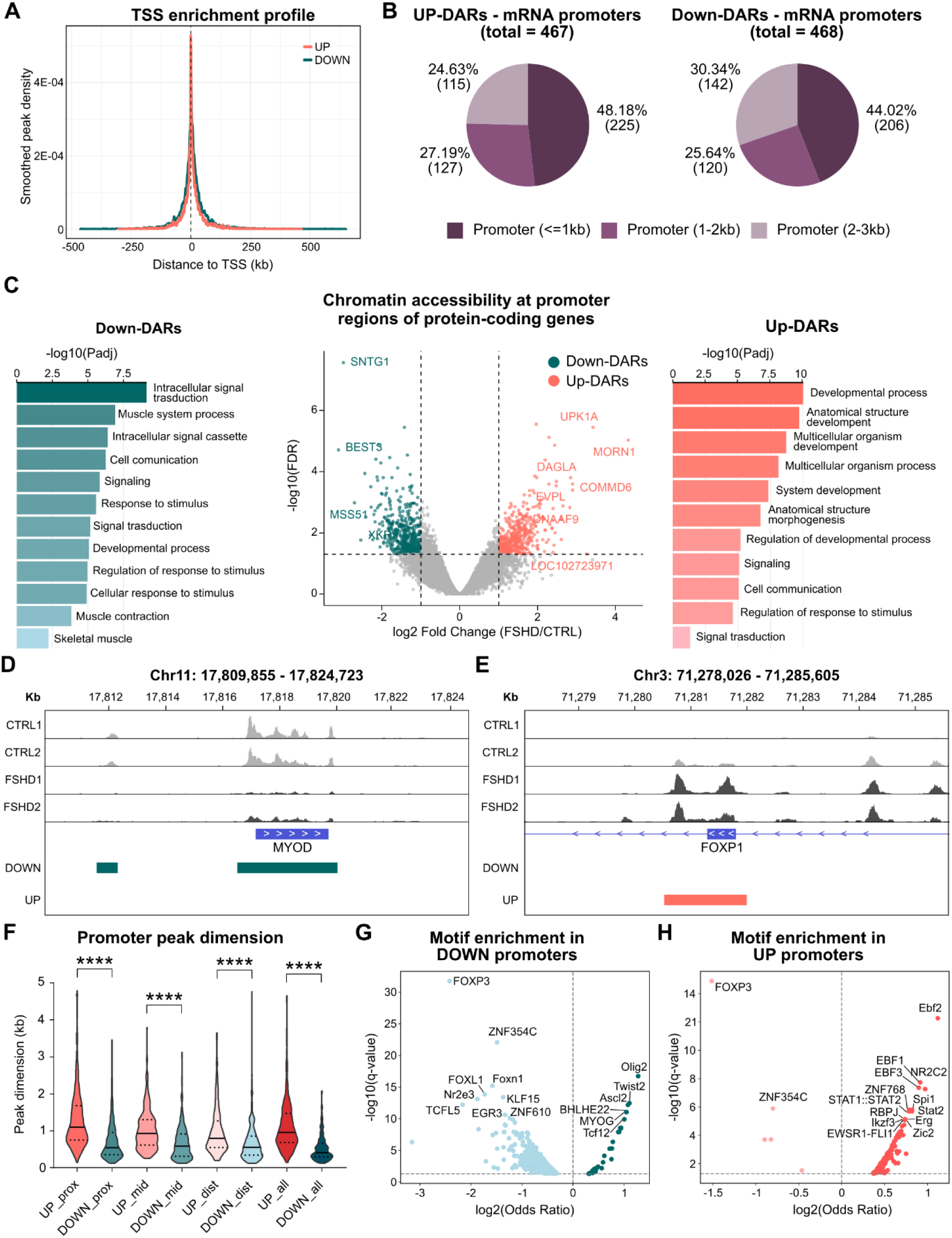
Chromatin accessibility and regulatory features at promoter regions of protein-coding genes in FSHD myoblasts. (A) TSS enrichment profile showing the distribution of ATAC-seq signal centered on annotated transcription start sites (TSS). The x-axis indicates the distance from the TSS (kb), and the y-axis shows the smoothed peak density. Profiles are shown separately for regions with increased accessibility in FSHD (Up-DARs, red) and decreased accessibility in FSHD (Down-DARs, teal). (B) Distribution of differentially accessible regions overlapping promoter regions of protein-coding genes. Promoter-associated DARs were categorized based on their distance from the TSS: proximal promoter (≤1 kb), intermediate promoter (1–2 kb), and distal promoter (2–3 kb). Pie charts show the proportion and absolute counts of Up-DARs (total = 467) and Down-DARs (total = 468) in each promoter interval. (C) Volcano plot showing chromatin accessibility changes at promoter regions of protein-coding genes. Each point represents an individual promoter-associated ATAC-seq peak. The x-axis indicates log₂ fold change (FSHD/CTRL), and the y-axis shows −log₁₀(FDR). Up-DARs are shown in red and Down-DARs in teal. Selected genes are labeled. Bar plots on the left and right show selected Gene Ontology (GO) biological process terms associated with genes linked to Down-DARs and Up-DARs, respectively. The x-axis indicates −log₁₀(adjusted P-value). (D) Representative Integrative Genomics Viewer (IGV) screenshot showing an example of a DOWN-DAR overlapping a promoter region of a protein-coding gene. ATAC-seq signal tracks (scale 0-150) from control and FSHD samples are shown aligned to genomic coordinates. (E) Representative IGV screenshot showing an example of a UP-DAR overlapping a promoter region of a protein-coding gene (scale 0-130), displayed as described in panel D. (F) Distribution of peak width for promoter-associated DARs. Violin plots show peak dimension (kb) for Up-DARs and Down-DARs stratified by promoter distance category (proximal, intermediate, distal) and combined categories. Individual data points and median values are shown. Statistical significance is indicated (****). (G) Motif enrichment analysis for transcription factor binding motifs within promoter regions associated with Down-DARs. Each point represents a transcription factor motif. The x-axis shows log₂ odds ratio, and the y-axis shows −log₁₀(q-value). Selected transcription factors are labeled. (H) Motif enrichment analysis for transcription factor binding motifs within promoter regions associated with Up-DARs. Each point represents a transcription factor motif. The x-axis shows log₂ odds ratio, and the y-axis shows −log₁₀(q-value). Selected transcription factors are labeled.

Promoters showing increased accessibility in FSHD were strongly enriched for Gene Ontology categories related to multicellular organism development, anatomical structure morphogenesis, regulation of developmental processes, cell adhesion, and cell fate specification (Figure 3C). This enrichment profile indicates aberrant acquisition of developmental and regulatory competence in proliferating myoblasts. In contrast, promoters exhibiting reduced accessibility were selectively associated with muscle-specific biological processes, including muscle contraction, intracellular signal transduction, sarcoplasmic reticulum calcium ion transport, and protein phosphorylation (Figure 3C), indicating coordinated loss of accessibility at loci central to myogenic identity and functional specialization (Figure 3C).

To illustrate these regulatory shifts, representative loci were selected for visualization by IGV based on functional relevance rather than peak magnitude alone. Promoters gaining accessibility included developmental regulators such as *EBF2*, *TRPC6*, and *FOXP1*, which displayed reproducible increases in accessibility across biological replicates (Figure 3E). At these loci, ATAC-seq signal extended beyond the core promoter region, generating broader accessible domains relative to controls. Conversely, promoters exhibiting reduced accessibility included key myogenic regulators such as *MYOD1*, *TTN*, and *RYR1*, which showed sharply reduced accessibility confined to the core promoter region in FSHD myoblasts (Figure 3D and Supplementary_Figure_S2).

Genome-wide quantitative analysis revealed differences in promoter accessibility architecture. While distributions of log fold change and distance from the TSS were comparable between Up- and Down-regulated promoter classes (Figure 3A,B), promoters gaining accessibility displayed significantly broader ATAC-seq peaks than promoters losing accessibility (mean peak width: 1279 bp vs. 894 bp). This pattern was consistently observed across proximal (1497 bp vs. 941 bp), intermediate (1119 bp vs. 752 bp), and distal promoter intervals (993 bp vs. 781 bp), indicating that accessibility gains are associated with spatial expansion of regulatory accessibility domains, whereas accessibility losses reflect localized erosion of promoter accessibility (Figure 3F).

To infer upstream regulatory mechanisms, motif enrichment analysis was performed on differentially accessible promoters relative to the full set of annotated promoters (Supplementary_Table_S2; Figure 3G,H). Promoters losing accessibility were significantly enriched for canonical myogenic bHLH and E-box motifs, including MYOD1, MYOG, MYF5/MYF6, and TCF-family factors (odds ratio ∼1.3–2.4; q-values down to ∼10^-17^), consistent with coordinated loss of accessibility at promoters embedded in the myogenic regulatory network. In contrast, promoters gaining accessibility displayed enrichment for motifs associated with alternative developmental and stress-responsive regulators, including EBF family members, STAT complexes, SPI1, and ETS-family transcription factors, indicating acquisition of non-myogenic regulatory competence.

These promoter-scale shifts demonstrate that promoter accessibility remodeling in FSHD myoblasts is highly directional and functionally coherent, involving coordinated loss of accessibility at promoters governing muscle identity and expansion of accessibility at promoters associated with developmental and stress-responsive regulatory programs.

### Chromatin accessibility losses cluster within chromosome scale architectural domains

To investigate whether chromatin accessibility changes in FSHD myoblasts display higher-order genomic organization, we analyzed the distribution of differentially accessible regions (DARs) at both chromosome and sub-chromosomal scales. For each autosome, the observed number of Up- and Down-regulated DARs was compared to the expected count based on genome-wide ATAC-seq peak density using a hypergeometric framework.

At the chromosome level, accessibility gains were broadly distributed and largely proportional to baseline chromatin accessibility. In contrast, accessibility losses exhibited pronounced chromosome-specific enrichment (Figure 4A,B and Supplementary_Table_S3). Significant over-representation of Down-DARs was detected on chromosomes 4 (observed = 434, expected = 369; FDR = 3.5 × 10⁻³), 13 (260 vs. 200; FDR = 4.0 × 10⁻⁴), and 18 (196 vs. 163; FDR = 3.5 × 10⁻²). Conversely, chromosome 19 showed marked depletion of Down-DARs, consistent with its high gene density and constitutively open chromatin state (Figure 4A). Up-DARs displayed limited chromosome-scale enrichment, with significant over-representation detected only on chromosome 8 (307 vs. 240; FDR = 1.9 × 10⁻⁴), indicating that accessibility gains are more diffusely distributed than accessibility losses (Figure 4B).

**Figure 4.**
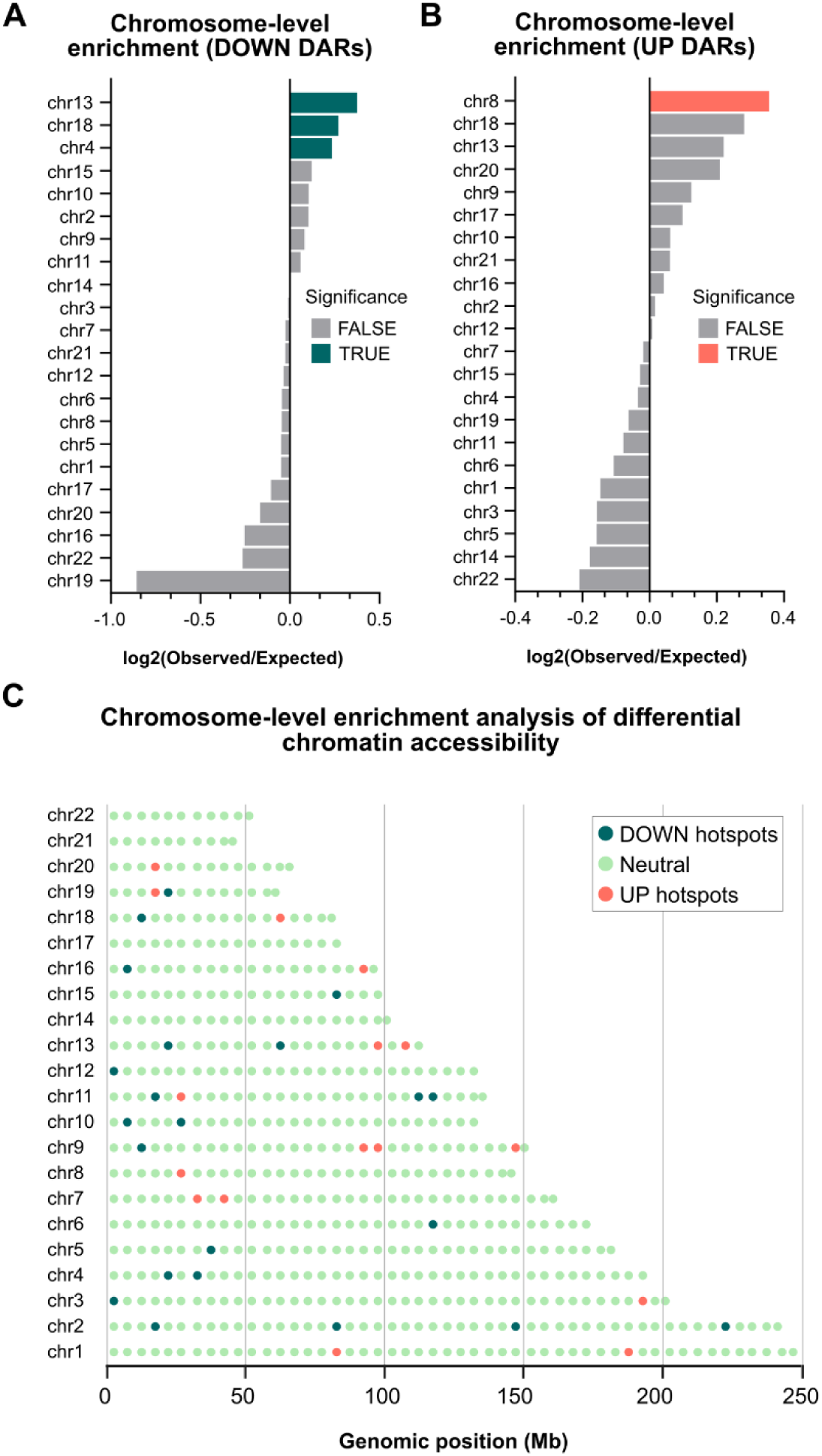
Chromosome-level enrichment and genomic distribution of differentially accessible chromatin regions in FSHD myoblasts. (A) Chromosome-level enrichment analysis of Down-DARs. Horizontal bar plot showing the log₂ ratio of observed versus expected numbers of Down-DARs for each human autosome. Each bar represents a chromosome. Values greater than zero indicate higher observed counts relative to expectation, and values less than zero indicate lower observed counts relative to expectation. Bars are colored according to statistical significance, as indicated in the legend. (B) Chromosome-level enrichment analysis of Up-DARs. Horizontal bar plot showing the log₂ ratio of observed versus expected numbers of Up-DARs for each chromosome. Chromosomes are ranked according to enrichment values. Bars are colored according to statistical significance, as indicated in the legend. (C) Genome-wide distribution of differential chromatin accessibility hotspots across chromosomes. Each horizontal row corresponds to an individual chromosome, arranged from chromosome 1 to chromosome 22. The x-axis indicates genomic position in megabases (Mb). Colored dots represent genomic bins classified according to enrichment status: Down-DAR hotspots (teal), Up-DAR hotspots (red), and neutral regions (green). Each dot corresponds to a genomic interval positioned according to its chromosomal coordinate.

To determine whether chromosome-level enrichment reflected diffuse effects or localized clustering, we performed a window-based enrichment analysis using non-overlapping 5-Mb intervals across the T2T-CHM13 genome as shown in (Figure 4C and Supplementary_Table_S3). For each window, observed and expected numbers of Up- and Down-regulated DARs were compared using hypergeometric tests followed by Benjamini-Hochberg correction. Among more than 600 analyzed windows, only a limited subset reached statistical significance (FDR < 0.05), indicating that accessibility remodeling is spatially constrained.

Significantly enriched windows were detected across multiple chromosomes and did not form a single continuous domain. Instead, enriched intervals appeared as isolated hotspots embedded within largely neutral chromatin landscapes (Figure 4C). This pattern indicates that accessibility changes are not uniformly distributed, but also do not propagate broadly along entire chromosome arms.

Together, these results indicate that chromatin accessibility remodeling in FSHD myoblasts is spatially restricted and organized into isolated genomic domains, partially overlapping with chromosomes exhibiting global enrichment, but also dispersed across the genome. This organization is consistent with regionally constrained perturbation of higher-order chromatin architecture rather than widespread chromosomal instability.

### Chromatin accessibility remodeling preferentially targets repressive nuclear compartments

To determine whether chromatin accessibility changes preferentially associate with specific nuclear compartments or chromatin states, we intersected differentially accessible regions (DARs) with published genome-wide maps of nucleolus-associated domains (NADs) (Peng et al. 2023) lamina-associated domains (LADs) derived from human fibroblast DamID datasets (NKI, UCSC Genome Browser)(Guelen et al. 2008), and chromatin state annotations based on ChromHMM from the ENCODE consortium (Ernst and Kellis 2012).

Because equivalent genome-wide maps are not available for primary human myoblasts, these datasets were used as reference maps. These reference maps capture major features of repressive genome compartmentalization that are broadly shared across human cell types, while allowing for cell type–specific differences in positioning and domain boundaries(Kumar et al. 2024). The results of these analyses are summarized in Supplementary_Table_S4-6 and Supplementary_Figure_S3.

Down-DARs showed significant enrichment in both NADs (OR = 1.10, 95% CI = 1.01–1.20, FDR = 2.6 × 10⁻⁴) and LADs (OR = 1.26, 95% CI = 1.18–1.35, FDR = 1.4 × 10⁻⁵) as shown in Figure 5A-D and Supplementary_Table_S6, indicating preferential localization of accessibility losses within repressive nuclear compartments. In contrast, Up-DARs were significantly depleted from NADs (OR = 0.86, FDR = 4.0 × 10⁻³) and LADs (OR = 0.68, FDR = 1.9 × 10⁻¹⁴).

**Figure 5.**
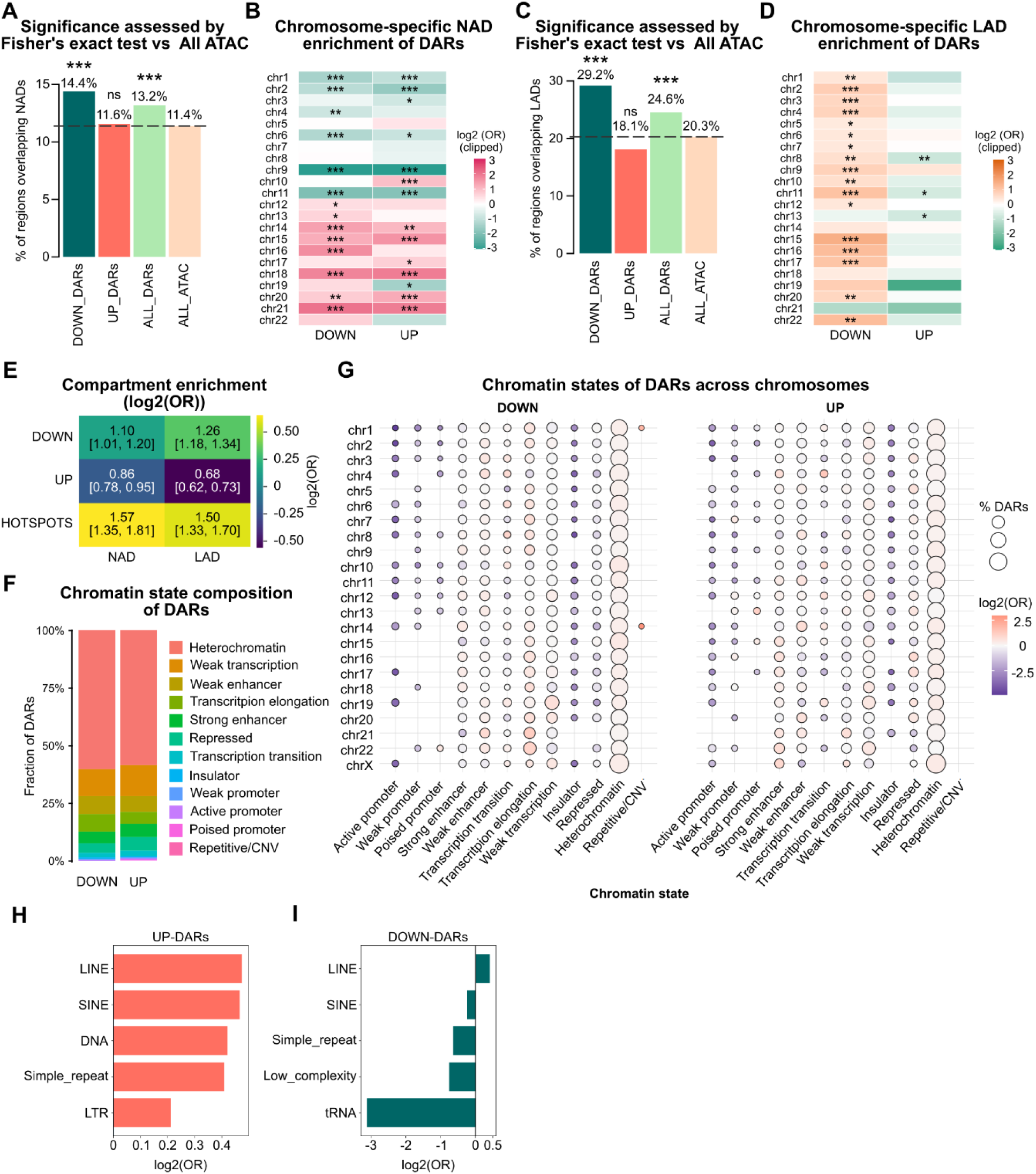
Association of differentially accessible chromatin regions with nuclear compartments, chromatin states, and repeat elements. (A) Percentage of Down-DARs, Up-DARs, all DARs, and all ATAC-seq peaks overlapping nucleolus-associated domains (NADs). Bar plots show the fraction of regions overlapping NADs. Statistical significance was assessed using Fisher’s exact test relative to all ATAC-seq peaks, as indicated. (B) Chromosome-specific enrichment of DARs within nucleolus-associated domains (NADs). Heatmap showing log₂ odds ratio (OR) values for enrichment of Down-DARs and Up-DARs across chromosomes 1–22. Each row represents a chromosome, and columns indicate Down-DARs and Up-DARs. Color intensity corresponds to log₂(OR), and statistical significance is indicated. (C) Percentage of Down-DARs, Up-DARs, all DARs, and all ATAC-seq peaks overlapping lamina-associated domains (LADs). Bar plots show the fraction of regions overlapping LADs. Statistical significance was assessed using Fisher’s exact test relative to all ATAC-seq peaks, as indicated. (D) Chromosome-specific enrichment of DARs within lamina-associated domains (LADs). Heatmap showing log₂ odds ratio values for enrichment of Down-DARs and Up-DARs across chromosomes. Each row corresponds to a chromosome, and columns indicate Down-DARs and Up-DARs. Color intensity represents log₂(OR), and statistical significance is indicated. (E) Compartment enrichment analysis showing log₂ odds ratio values for enrichment of Down-DARs, Up-DARs, and DAR hotspots in NAD and LAD compartments. Values are shown together with corresponding confidence intervals. (F) Chromatin state composition of DARs. Stacked bar plots showing the fraction of Down-DARs and Up-DARs assigned to different chromatin states, including heterochromatin, weak transcription, weak enhancer, transcription elongation, strong enhancer, repressed, transcription transition, insulator, weak promoter, active promoter, poised promoter, and repetitive/CNV. (G) Chromatin state distribution of DARs across chromosomes. Bubble plots showing the percentage of DARs associated with each chromatin state across chromosomes 1–22 and chromosome X. Circle size represents the fraction of DARs, and color indicates log₂ odds ratio for enrichment. Separate panels are shown for Down-DARs and Up-DARs. (H) Enrichment of repeat element classes in Up-DARs. Bar plot showing log₂ odds ratio values for overlap between Up-DARs and annotated repeat classes, including LINE, SINE, DNA elements, simple repeats, and LTR elements. (I) Enrichment of repeat element classes in Down-DARs. Bar plot showing log₂ odds ratio values for overlap between Down-DARs and annotated repeat classes, including LINE, SINE, simple repeats, low complexity regions, and tRNA elements.

Remodeling hotspots showed the strongest compartmentalization, with significant enrichment in both NADs (OR = 1.57, FDR = 1.2 × 10⁻²) and LADs (OR = 1.50, FDR = 2.7 × 10⁻⁴) (Figure 5E, Supplementary_Figure_S3 and Supplementary_Table_S6), indicating that clustered chromatin remodeling preferentially targets structurally constrained nuclear environments.

Analysis of significantly enriched chromosomes (Figure 5B,D and Table 2) revealed a reproducible segregation into distinct architectural classes. NAD-associated remodeling was restricted to a limited subset of chromosomes, including chromosomes 14, 15, 16, 18, and 21, characterized by large effect sizes, whereas LAD-associated remodeling involved a broader set of chromosomes and prominently included chromosome 4, which showed consistent enrichment within lamina-associated domains. In contrast, gene-dense chromosomes such as chromosomes 1, 2, 6, 9, and 11 were consistently depleted. Notably, chromosome 9 showed near-complete exclusion from nucleolus-associated remodeling, indicating selective compartmentalization of accessibility changes.

**Table 2:**
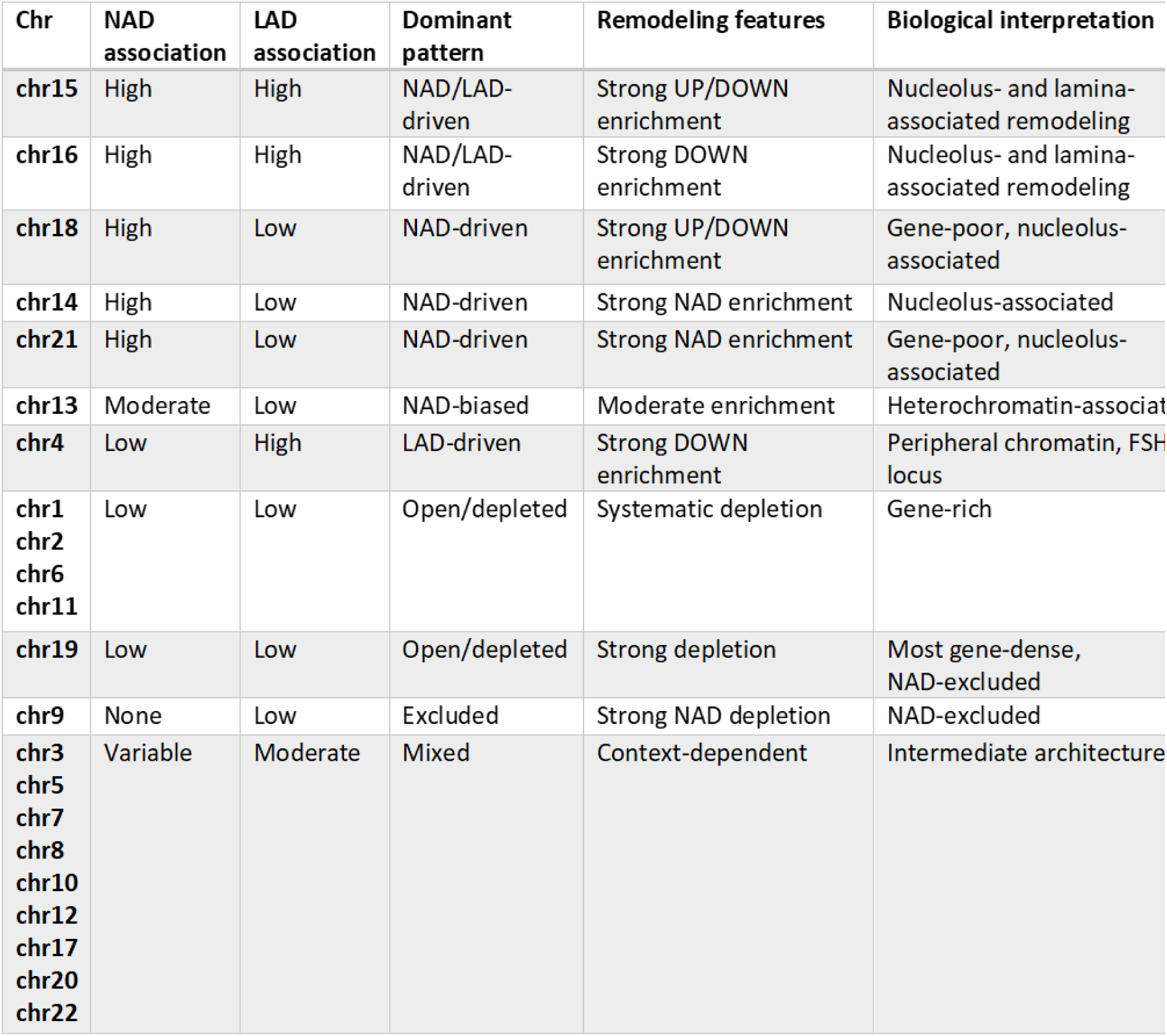
Chromosome-level architectural association of chromatin accessibility remodeling in FSHD myoblasts. Chromosomes were classified based on enrichment or depletion of differentially accessible regions within nucleolus-associated domains (NADs) and lamina-associated domains (LADs). Classification is based on odds ratio and FDR-corrected enrichment analysis. Chromosomes segregate into distinct architectural classes including NAD/LAD-driven, NAD-driven, LAD-driven, open/depleted, excluded, and mixed patterns, reflecting differential involvement of nuclear compartments in chromatin accessibility remodeling.

| Chr | NAD association | LAD association | Dominant pattern | Remodeling features | Biological interpretation |
| --- | --- | --- | --- | --- | --- |
| chr15 | High | High | NAD/LAD-driven | Strong UP/DOWN enrichment | Nucleolus- and lamina-associated remodeling |
| chr16 | High | High | NAD/LAD-driven | Strong DOWN enrichment | Nucleolus- and lamina-associated remodeling |
| chr18 | High | Low | NAD-driven | Strong UP/DOWN enrichment | Gene-poor, nucleolus-associated |
| chr14 | High | Low | NAD-driven | Strong NAD enrichment | Nucleolus-associated |
| chr21 | High | Low | NAD-driven | Strong NAD enrichment | Gene-poor, nucleolus-associated |
| chr13 | Moderate | Low | NAD-biased | Moderate enrichment | Heterochromatin-associated |
| chr4 | Low | High | LAD-driven | Strong DOWN enrichment | Peripheral chromatin, FSH locus |
| chr1<br>chr2<br>chr6<br>chr11 | Low | Low | Open/depleted | Systematic depletion | Gene-rich |
| chr19 | Low | Low | Open/depleted | Strong depletion | Most gene-dense, NAD-excluded |
| chr9 | None | Low | Excluded | Strong NAD depletion | NAD-excluded |
| chr3<br>chr5<br>chr7<br>chr8<br>chr10<br>chr12<br>chr17<br>chr20<br>chr22 | Variable | Moderate | Mixed | Context-dependent | Intermediate architecture |

To define the epigenomic context of these accessibility changes, we performed genome-wide chromatin state segmentation using ChromHMM by integrating ATAC-seq with multiple histone modification profiles (Supplementary_Table_S7 and Figure 5F). This hidden Markov model–based approach identified recurrent regulatory and structural chromatin states, including active promoters, enhancers, weakly transcribed regions, and repressed or heterochromatic domains.

Mapping DARs onto chromatin state annotations revealed a strongly non-random distribution. Both Up- and Down-DARs were significantly depleted from active promoter states and enriched in weakly transcribed and repressed chromatin, indicating preferential accessibility remodeling outside canonical regulatory promoters (Figure 5F). State transition analysis further showed that Down-DARs frequently corresponded to shifts from active or poised configurations toward repressed chromatin, whereas Up-DARs were associated with transitions toward more permissive environments. Enrichment analysis revealed preferential localization of DARs within heterochromatin-associated, low-signal, and weakly transcribed chromatin (OR ≈ 1.1–1.3) (Supplementary_Table_S7), accompanied by marked depletion from promoter and insulator regions. Chromosome-resolved analyses showed that this distribution extends across multiple chromosomes, indicating that accessibility remodeling reflects a genome-wide architectural pattern rather than promoter-centered dysregulation (Figure 5G).

Consistent with these observations, analysis of RepeatMasker annotations revealed distinct repeat class associations for accessibility gains and losses (Supplementary_Table_S8). Figure 5H,I shows that Up-DARs were significantly enriched in multiple retrotransposon-derived elements, including SINEs, LINEs, DNA transposons, simple repeats, and LTR elements (FDR < 0.05), indicating preferential accessibility gains within repeat-rich genomic regions. In contrast, Down-DARs showed significant depletion of several repeat classes associated with accessible retrotransposon-derived chromatin, including SINEs, simple repeats, and low-complexity elements. This indicates that accessibility losses preferentially occur within structurally constrained heterochromatin domains that are not characterized by retrotransposon-associated accessible chromatin. Instead, they represent stabilized architectural regions of the nuclear periphery and nucleolar compartment.

The same directional signal recurs across independent annotations: accessibility losses concentrate in NAD/LAD-linked and heterochromatin/low-signal states, while gains favor repeat-associated permissive contexts. This convergence supports an architectural redistribution of accessibility rather than just promoter-centered dysregulation.

### Regional chromatin remodeling at 4q35 is uncoupled from accessibility at canonical DUX4 target genes

Given the central involvement of chromosome 4q35 in FSHD pathogenesis, we next examined chromatin accessibility changes along the distal region of chromosome 4 at higher resolution. Notably, despite the improved resolution afforded by the T2T assembly, no reproducible ATAC-seq peaks were detected within the D4Z4 repeat array itself in either FSHD or control samples. Across all biological replicates, this region remained largely inaccessible, and no consistent disease-associated differences were observed (Supplementary_Figure_S4A). This indicates that, at the level of bulk chromatin accessibility, D4Z4 remains refractory to transposase integration in proliferating myoblasts, even in the disease context. Consistently, no reproducible disease-associated opening was detected within or distal to the D4Z4 array, and accessibility signals declined beyond the DUX4L locus, limiting coverage of FRG2 and adjacent repeat-derived transcripts.

Sliding window analysis of differentially accessible regions (DARs) revealed a non-uniform distribution of accessibility losses (Supplementary_Table_S9 and Supplementary_Figure_S4B), with several megabase-scale intervals displaying enrichment of Down-DARs relative to the chromosome-wide average. Analysis of the distal 4q35 interval (chr4:188 Mb to telomere) identified a limited and heterogeneous set of accessibility changes (22 Up- and 29 Down-DARs), arguing against a model of uniform chromatin opening. Instead, remodeling at this locus was spatially confined and bidirectional, consistent with localized reorganization of higher-order chromatin structure rather than passive derepression of the repeat array.

Most reproducible ATAC-seq peaks in this region were restricted to a limited number of promoter-associated elements located several megabases centromeric to the D4Z4 array, including those linked to *TENM3*, WWC2-AS, *MYL12BP2*, and *SLC25A4*. These promoters represent the main accessible regulatory elements detectable by short-read ATAC-seq within the distal 4q35 interval, even though they are spatially distant from the core pathogenic locus. To assess whether these accessibility patterns were associated with changes in transcriptional output, we performed quantitative RT-qPCR on the same samples used for ATAC-seq analysis, measuring transcript levels of multiple genes distributed along the 4q35 locus and normalizing to GAPDH mRNA (Supplementary_Figure_S4C). For most genes, expression levels were comparable between control and FSHD myoblasts, consistent with the largely stable accessibility profiles observed by ATAC-seq. Notably, the only transcripts showing reproducible and significant differences between control and FSHD samples were located in the telomeric portion of the locus, in agreement with our previous reports (Salsi et al. 2025b). These transcripts have been shown to be regulated not only at the transcriptional level but also through post-transcriptional mechanisms, indicating that their deregulation cannot be inferred solely from chromatin accessibility measurement.

Given the lack of overt accessibility changes at the D4Z4 locus, the limited involvement of nearby promoters, and the largely stable transcriptional output across 4q35, we next asked whether chromatin accessibility remodeling reflected activation of the canonical DUX4-driven transcriptional program. To this end, we analyzed ATAC-seq profiles across annotated DUX4-responsive loci (Geng et al. 2012; Young et al. 2013; Hendrickson et al. 2017; Jiang et al. 2020; Bosnakovski et al. 2023). (Supplementary_Table_S9), focusing on regions spanning ±2 kb from transcription start sites to transcription end sites.

Aggregate accessibility profiles revealed comparable promoter-centered patterns in control and FSHD samples, without the emergence of disease-specific accessibility gains. At the single-gene level, accessibility was highly heterogeneous and not concordant across FSHD replicates. Accordingly, overlap between DARs and DUX4 target loci was minimal, with only two targets intersecting Up-DARs and none overlapping Down-DARs. These observations indicate that chromatin accessibility changes captured by ATAC-seq are largely uncoupled from the canonical DUX4 transcriptional program in proliferating myoblasts.

Importantly, the core pathogenic region at 4q35 resides within highly repetitive and low-mappability sequences that are intrinsically refractory to short-read accessibility assays. Therefore, the observed remodeling patterns most likely reflect indirect or flanking chromatin responses associated with locus deregulation and nuclear repositioning, rather than direct opening of the D4Z4 array.

These data support a model in which FSHD-associated chromatin remodeling at 4q35 is spatially restricted, architecturally complex, and primarily involves surrounding heterochromatin-rich domains rather than canonical regulatory elements

## Discussion

### Stage-dependent chromatin accessibility remodeling in FSHD myoblasts

This study demonstrates that chromatin accessibility remodeling in facioscapulohumeral muscular dystrophy emerges during the proliferative myoblast stage and preferentially affects non-coding and structurally constrained genomic domains. Rather than solely reflecting focal dysregulation of muscle gene promoters, accessibility changes are distributed at chromosome and domain scales, indicating a global redistribution of regulatory accessibility. These findings suggest that chromatin dysregulation in FSHD reflects early perturbation of nuclear regulatory architecture.

Importantly, these accessibility differences are largely attenuated upon terminal differentiation, indicating that architectural remodeling defines a stage-specific vulnerability of proliferating myoblasts rather than a persistent feature of differentiated muscle. This observation is consistent with previous studies demonstrating that FSHD myoblasts represent a uniquely sensitive regulatory state, in which genome organization, transcriptional competence, and translational control remain highly plastic and responsive to perturbation (Salsi et al. 2025b, 2025a). In particular, quantitative proteomic analyses have revealed activation of compensatory translational programs in proliferating FSHD myoblasts that are not sustained upon differentiation, suggesting that early architectural and regulatory instability may precede and contribute to later cellular dysfunction (Salsi et al. 2025a). Clinical and experimental observations reinforce the relevance of this early window. Early-life physical activity has been associated with reduced disease severity (Bettio et al. 2024) and epigenetic pharmacological interventions have demonstrated measurable regulatory effects (Salsi et al. 2025b; Nordlinger et al. 2026), supporting the idea that FSHD chromatin states retain partial plasticity. Given the the importance of protein homeostasis maintenance in skeletal muscle and its progressive decline with aging (Sandri 2013; López-Otín et al. 2013; Salsi et al. 2023), early architectural instability may increase susceptibility to cumulative differentiation- and age-associated stress. These findings establish proliferating myoblasts as a critical window of vulnerability in FSHD pathogenesis, during which disruption of chromatin organization and regulatory competence is established prior to terminal differentiation and downstream gene expression abnormalities.

### Chromosome-scale remodeling and uncoupling from canonical DUX4-driven transcription

At the chromosome scale, chromatin accessibility changes in FSHD myoblasts exhibit pronounced higher-order organization, with accessibility losses clustering within discrete megabase-scale domains, in agreement with earlier reports of large-scale chromatin and nuclear reorganization in FSHD (Bodega et al. 2009; Broucqsault et al. 2013; Robin et al. 2014; Gaillard et al. 2019). Given the central role of the 4q35 region in FSHD, we examined the distal portion of chromosome 4 at higher resolution. However, despite the use of a complete genome assembly, we detect no reproducible accessibility within the D4Z4 repeats themselves, consistent with previous observations that this region remains embedded in repressive chromatin environments (Lyle et al. 1995; Masny et al. 2004; Tam et al. 2004; Ottaviani et al. 2009).

These findings support regulatory mechanisms based on long-range interactions and nuclear repositioning, in contrast with simplified derepression models (Snider et al. 2010). Indeed, accessibility changes show limited correspondence with steady-state transcriptional output across the locus, and only telomeric transcripts, previously shown to be subject to multilayered transcriptional and post-transcriptional regulation (Salsi et al. 2025b), display reproducible deregulation. Moreover, analysis of canonical DUX4 target loci reveals no coherent accessibility signature in proliferating myoblasts, extending prior work showing that DUX4 activity can be transient, heterogeneous, and uncoupled from stable epigenomic states (Snider et al. 2010; Tassin et al. 2013; Haynes et al. 2018) ATAC-seq studies performed in experimental systems with induced or high-level DUX4 expression have demonstrated robust chromatin opening at canonical DUX4 target loci, consistent with pioneer-like activity of DUX4 (Vuoristo et al. 2022; Bosnakovski et al. 2023; Young et al. 2013). However, these models rely on synchronized and elevated DUX4 activity that does not fully recapitulate the endogenous FSHD context. Instead, in primary FSHD myoblasts, where DUX4 expression is rare and heterogeneous, accessibility changes at canonical DUX4 targets are limited and not reproducibly detected at the population level. These observations indicate that early chromatin remodeling in FSHD reflects higher-order architectural perturbations rather than a direct and stable chromatin accessibility footprint of DUX4 activity.

### Architectural and regulatory layers of chromatin dysregulation in FSHD

Chromatin accessibility remodeling in FSHD preferentially targets genomic domains associated with higher-order nuclear organization, particularly heterochromatin-enriched and nucleolus-associated compartments. Because genome-wide maps of nucleolus-associated and lamina-associated domains are not available for primary human myoblasts, we relied on reference datasets generated in other human cell types. Although the precise positioning of individual domains can vary between cell types, heterochromatin-associated compartments represent a fundamental structural component of nuclear genome organization (Shah et al. 2023; Kumar et al. 2024). Importantly, the accessibility changes identified here occur at megabase-scale domains and show consistent enrichment across multiple independent compartment annotations and chromatin state classifications, supporting a robust association between chromatin accessibility loss and heterochromatin-associated nuclear architecture.

At the megabase scale, accessibility losses cluster within discrete genomic regions enriched for nucleolus-associated domains, which correspond to late-replicating, repeat-rich, and structurally constrained heterochromatic regions (Vertii et al. 2019; Bersaglieri et al. 2022; Peng et al. 2023; Kumar et al. 2024). This pattern indicates that chromatin accessibility remodeling in FSHD preferentially affects genomic compartments linked to nucleolar organization and higher-order genome topology, implicating nuclear architecture as a major substrate of epigenomic dysregulation. Consistent with this architectural specificity, repeat class analysis reveals distinct distributions of accessibility gains and losses across repetitive elements. Accessibility gains preferentially occur within retrotransposon-associated sequences, including SINE and LINE elements, whereas accessibility losses preferentially localize within structurally constrained heterochromatin domains enriched in nucleolus- and lamina-associated compartments but depleted in repeat classes associated with accessible chromatin. This selective redistribution of accessibility indicates that chromatin remodeling reflects compartment-specific architectural perturbations rather than uniform derepression of repetitive DNA.

At the promoter scale, accessibility remodeling follows a distinct regulatory pattern characterized by directional motif enrichment patterns. Promoters losing accessibility are enriched for canonical myogenic regulatory motifs, including E-box and bHLH-associated elements, consistent with reduced accessibility of core myogenic regulatory regions. In contrast, promoters gaining accessibility preferentially harbor motifs associated with alternative developmental and stress-responsive regulatory programs, indicating redistribution of regulatory competence toward non-myogenic transcriptional inputs. Because ATAC-seq measures transcription factor-permissive chromatin rather than transcription itself, these motif signatures reflect altered regulatory accessibility rather than direct transcriptional activation. Together, these findings indicate that chromatin configuration in FSHD operates across multiple hierarchical levels of genome organization. Megabase-scale remodeling affects heterochromatin-associated nuclear compartments and genome topology, whereas promoter accessibility changes reflect localized alterations in regulatory competence. This multilayered organization of chromatin dysregulation is summarized schematically in Figure 6.

**Figure 6.**
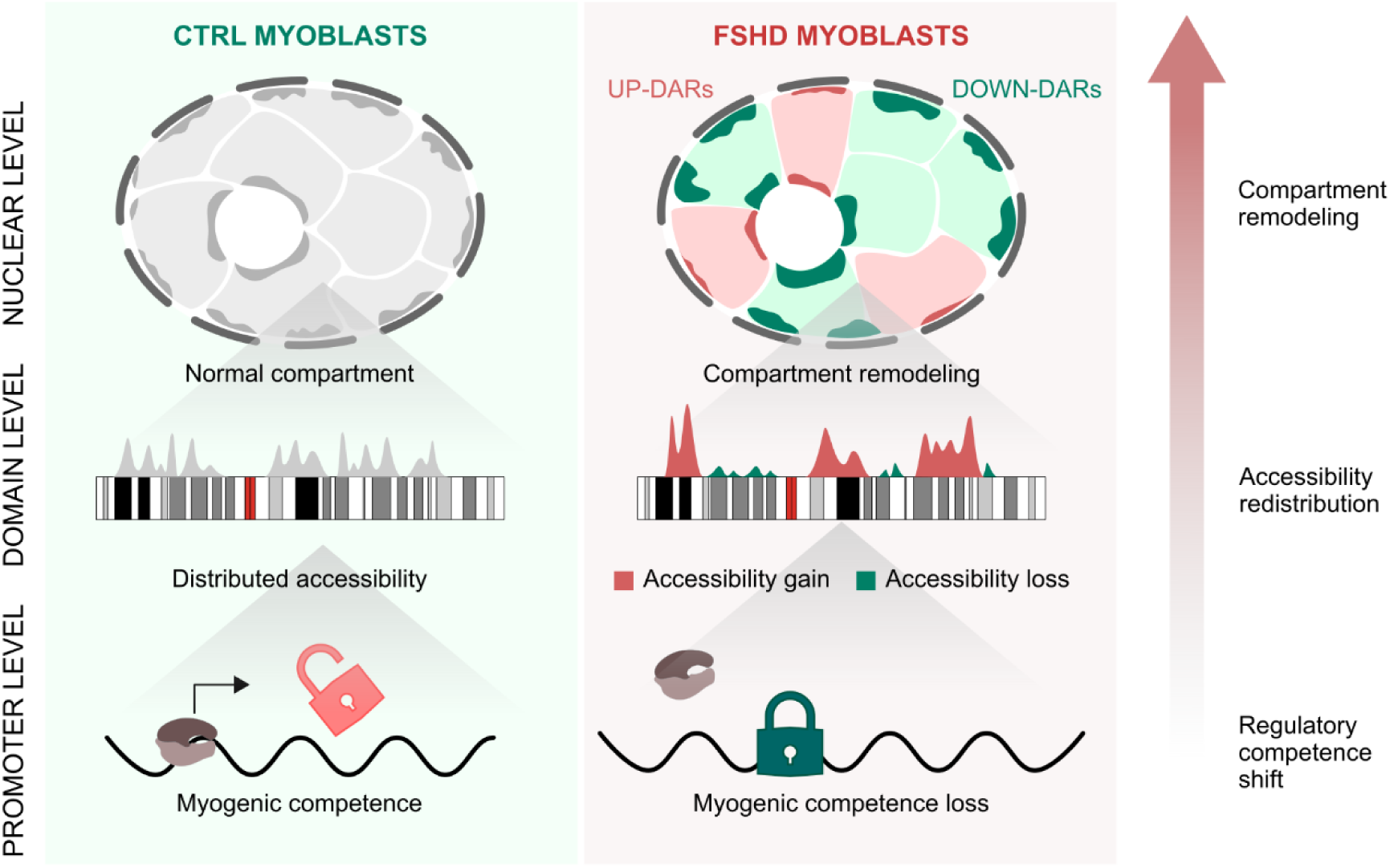
Conceptual model of chromatin accessibility remodeling in FSHD myoblasts. Schematic model illustrating the multilayered consequences of chromatin accessibility remodeling in FSHD myoblasts. In control myoblasts (left), higher-order nuclear architecture maintains stable chromatin compartmentalization, resulting in balanced chromatin accessibility across genomic domains and preservation of myogenic regulatory competence. In FSHD myoblasts (right), chromatin accessibility remodeling occurs across multiple regulatory layers. At the nuclear level, chromatin compartments undergo structural reorganization. At the domain level, accessibility is redistributed across genomic regions exhibiting accessibility gains (Up-DARs) and losses (Down-DARs). At the promoter level, these changes translate into altered regulatory competence affecting gene expression programs. Together, these layers illustrate how architectural perturbations propagate from higher-order genome organization to local regulatory elements in FSHD myoblasts.

### Architectural remodeling implicates nucleolar-associated genome organization and repeat-derived RNA regulatory mechanisms

The enrichment of accessibility loss within NAD- and LAD-linked domains indicates that FSHD myoblasts exhibit a compartment-focused collapse of regulatory accessibility across structurally constrained heterochromatin. This pattern does not demonstrate increased physical clustering per se, but it supports an architectural model in which early disease-associated remodeling is guided by nuclear compartment topology, consistent with RNA-mediated nucleolar genome organization pathways implicated in FSHD.

The preferential involvement of nucleolus-associated genomic domains is particularly notable in light of accumulating evidence that repeat-derived long non-coding RNAs participate in nucleolar genome organization. In particular, the repeat-derived lncRNA FRG2A, consistently overexpressed in FSHD muscle cells, localizes to nucleolar-associated chromatin and has been functionally linked to rDNA regulation and nucleolar activity. The overlap between accessibility loss domains and nucleolus-associated compartments is therefore consistent with engagement of nucleolus-linked regulatory networks in the architectural chromatin accessibility state observed in FSHD myoblasts. Supporting this view, enrichment analysis of promoter-associated non-coding DARs reveals significant associations with RNA processing and nucleolar functions, indicating that accessibility remodeling intersects pathways connected to nucleolar genome regulation.

More broadly, repeat-derived lncRNAs are increasingly recognized as key regulators of nuclear architecture, acting as molecular scaffolds that organize chromatin domains and nuclear compartments. Examples such as XIST, NEAT1, FIRRE, TERRA and FRG2A illustrate how non-coding RNAs mediate higher-order genome organization, with major consequences for gene regulation and cellular identity (Tinker and Brown 1998; West et al. 2014; Hacisuleyman et al. 2014; Chu et al. 2017; Salsi et al. 2025a).

These findings place FSHD within an emerging class of nuclear architecture disorders in which pathogenic effects lead to disruption of genome topology and nuclear compartment organization rather than isolated gene-level dysregulation. Similar architectural mechanisms have been implicated in laminopathies, progeroid syndromes, neurodegenerative disorders, and repeat expansion diseases (Taimen et al. 2009; Schreiber and Kennedy 2013; Zhang et al. 2015), where disruption of nuclear compartmentalization precedes overt transcriptional dysfunction.

We propose that disruption of RNA-mediated architectural networks in proliferating myoblasts contributes to redistribution of chromatin accessibility and altered regulatory competence, thereby increasing cellular vulnerability prior to overt transcriptional and differentiation defects. Rather than implying a direct causal mechanism, this architectural state defines a stage-specific chromatin configuration that emerges during myoblast proliferation and reflects impaired stabilization of higher-order genome organization. In this framework, RNA-mediated architectural interactions normally contribute to maintaining nuclear compartment integrity, whereas their perturbation in FSHD delineates a temporal window characterized by increased regulatory plasticity and susceptibility to differentiation and proteostasis demand. This architecture-centered perspective shifts attention toward upstream regulatory mechanisms that may influence disease susceptibility and progression, and provides a foundation for future studies aimed at identifying mechanistic and therapeutic targets.

##### Focus Box 1 Early architectural chromatin accessibility state in FSHD

- **Remodeling is strongly stage-dependent.**

In proliferating myoblasts, we detect 12,593 DARs (7,491 Down, 5,102 Up), whereas differentiated myotubes show <200 DARs with markedly reduced effect sizes, indicating that accessibility remodeling is largely restricted to the proliferative stage.

- **Stage-specific remodeling preferentially targets non-coding regulatory space.**

Across myoblast DARs, ∼70–80% map to intronic/distal intergenic regions, while only ∼13–17% overlap promoters (-3 kb from the TSS), indicating global redistribution of accessibility rather than promoter-centered dysregulation.

- **Promoter changes indicate a shift in regulatory competence**.

Promoters gaining accessibility are enriched for developmental and stress-responsive programs, whereas promoters losing accessibility map to muscle identity and intracellular signaling, consistent with erosion of a myogenic regulatory state.

- **Accessibility losses follow higher-order genomic organization**.

Down-DARs are non-uniformly distributed across chromosomes and cluster into discrete megabase-scale domains, revealing architectural patterning of chromatin remodeling.

- **4q35 undergoes regional remodeling without detectable D4Z4 opening**.

Accessibility changes occur in domains flanking the pathogenic locus, but the D4Z4 array remains largely inaccessible in bulk ATAC-seq even in the T2T-CHM13 assembly.

- **Genome-wide remodeling is largely uncoupled from a canonical DUX4 accessibility program.**

DUX4 target loci do not show a coherent or reproducible accessibility signature in myoblasts, indicating that widespread remodeling occurs largely independently of sustained DUX4-driven chromatin opening.

- **Overall, these findings define a stage-specific architectural accessibility state in FSHD. Accessibility is redistributed across repeat-rich and structurally constrained genomic domains before terminal differentiation, supporting a model in which nuclear architecture defects precede overt gene-centric regulatory reprogramming.**

## Materials and Methods

### Primary human myoblast cultures and myogenic differentiation

Primary human myoblast cultures were obtained from FSHD patients and unaffected controls belonging to well-characterized familial cohorts. Control and FSHD-derived primary myoblasts were selected from the Italian National Registry for FSHD (20), and molecular and clinical features are reported in Supplementary_Figure_S1. Myoblasts were purified from muscle biopsies (Supplementary_Table_S1) as previously described (Salsi et al. 2025b). Cells were expanded under proliferative conditions in growth medium optimized to preserve myogenic potential and maintained at subconfluent density. Briefly, cells were cultured in DMEM supplemented with 20% fetal bovine serum (FBS), 1% L-glutamine, 1% penicillin–streptomycin, 2 ng/mL epidermal growth factor (EGF), and 25 ng/mL fibroblast growth factor (FGF). Cells were detached using 0.25% trypsin diluted 1:10 in PBS.

Myogenic differentiation was induced by replacing growth medium with differentiation medium and maintaining cultures until formation of multinucleated myotubes, as previously described (Salsi et al. 2025b). Samples were collected at defined developmental stages corresponding to proliferating myoblasts and differentiated myotubes. All experiments were performed using independent biological replicates. Cells were harvested directly for ATAC-seq library preparation without fixation.

### ATAC-seq

ATAC-seq libraries were generated using the ATAC-Seq Kit (Active Motif, Cat. No. 53150) following the manufacturer’s protocol. For each biological replicate, 100,000 viable cells were collected, washed in ice-cold PBS, and lysed in ATAC Lysis Buffer to isolate intact nuclei. Nuclei were pelleted by centrifugation and immediately resuspended in Tagmentation Master Mix containing assembled Tn5 transposase, which simultaneously fragments accessible chromatin regions and inserts sequencing adapters.

The tagmentation reaction was carried out at 37 °C for 30 minutes with agitation. Following transposition, DNA fragments were purified using silica column purification and eluted in DNA Elution Buffer.

Purified tagmented DNA was amplified by PCR using indexed Nextera-compatible primers and Q5 High-Fidelity DNA Polymerase. Amplification conditions included an initial extension step at 72 °C, followed by denaturation at 98 °C and 10 amplification cycles.

Amplified libraries were purified using SPRI bead cleanup and eluted in Elution Buffer. Library concentration and size distribution were assessed prior to sequencing. Libraries were subjected to paired-end sequencing on an Illumina platform.

Library quality and fragment size distribution were assessed prior to sequencing. Paired-end sequencing (150 bp reads) was performed by Macrogen on an Illumina NovaSeq X platform, generating approximately 8 Gb of sequencing data per library.

### ATAC-seq read processing, alignment, and peak calling

Primary read processing and downstream ATAC-seq data analysis were performed by Macrogen (Seoul, South Korea). Adapter trimming and quality filtering were conducted using Trim Galore. Filtered reads were aligned to the telomere-to-telomere T2T-CHM13v2.0 reference genome using Bowtie2 (Langmead and Salzberg 2012). PCR duplicate reads, mitochondrial reads, and reads overlapping ENCODE blacklist regions were removed prior to peak calling. Accessible chromatin regions were identified genome-wide using MACS2 peak calling (Zhang et al. 2008).

Differential accessibility analysis between FSHD and control samples was performed using the csaw/edgeR statistical framework, which applies normalization, dispersion estimation, and quasi-likelihood testing to identify differentially accessible regions. P-values were adjusted using the Benjamini-Hochberg procedure to control the false discovery rate (FDR), and differentially accessible regions (DARs) were defined using FDR < 0.05 and an absolute log₂ fold change ≥ 1.

Standard ATAC-seq quality control metrics, including mapping rate, non-redundant fraction (NRF), transcription start site enrichment score (TSSES), and fraction of reads in peaks (FRiP), confirmed high-quality libraries suitable for downstream analysis.

### RNA extraction and Real-time quantitative PCR (RT-qPCR)

Total cellular RNA was obtained from cell lines and HPMs by using PureLink RNA Mini Kit (Thermo Fisher Scientific cat #12183018A), according to the manufacturer’s instructions. DNAse digestion and cDNA synthesis were performed by using Maxima H-cDNA Synthesis Master Mix, with dsDNase (Thermo Fisher M1482). Specific mRNA expression was assessed by qRTPCR (iTaq Universal SYBR® Green Supermix, BIORAD #1725120 in a CFX connect Real Time Machine BIORAD) using primers described in (Salsi et al. 2025b) normalized over GAPDH housekeeping mRNAs.

### Genome assembly and genomic coordinate processing

All genomic analyses were performed using the telomere-to-telomere T2T-CHM13 reference genome assembly to enable accurate representation of repeat-rich, subtelomeric, and structurally complex genomic regions that are incompletely resolved in previous genome builds. Where required for compatibility with external annotation datasets, genomic coordinates were converted between genome assemblies using the UCSC LiftOver tool (Hinrichs et al. 2006). Only genomic intervals successfully mapped to the T2T-CHM13 assembly were retained. Genomic interval manipulation, overlap analysis, and coordinate processing were performed using the GenomicRanges package in R(Lawrence et al. 2013).

### Functional enrichment analysis using g:Profiler

Functional enrichment analysis was performed using g:Profiler (g:GOSt)(Raudvere et al. 2019) on genes associated with promoter-overlapping differentially accessible regions (DARs). Promoters were defined as genomic intervals spanning -3 kb from annotated transcription start sites (TSS) (T2T-CHM13 annotation). Promoter-associated DARs were stratified by direction of change into Up-promoters and Down-promoters, and analyzed separately for (i) protein-coding genes and (ii) non-coding RNA genes.

Enrichment was assessed for Gene Ontology terms (biological process, molecular function, cellular component) and relevant pathway resources as implemented in g:Profiler. Multiple testing correction was performed using the g:Profiler default method, and terms passing the adjusted significance threshold (FDR < 0.05) were retained. Enrichment results were visualized by ranking terms by adjusted P-value and plotting −log10(adjusted P-value) for selected representative categories.

### Motif enrichment analysis at promoter regions

Motif enrichment analysis was performed on differentially accessible promoter regions identified by ATAC-seq. Promoters were defined as genomic intervals spanning 3 kb upstream of annotated transcription start sites (TSS) (−3 kb to 0) according to the T2T-CHM13 genome annotation. Differentially accessible promoters were classified into two groups according to ATAC-seq signal differences between FSHD and control myoblasts: promoters showing increased accessibility (Up-promoters) and promoters showing decreased accessibility (Down-promoters).

Motif enrichment was assessed using position weight matrices (PWMs) obtained from the JASPAR 2024 CORE vertebrate database(Castro-Mondragon et al. 2022). Motif occurrences were identified using the Bioconductor package motifmatchr (version 1.32.0), which scans genomic sequences for PWM matches using a statistical threshold corresponding to a motif match p-value of 1 × 10⁻⁴. For each motif, enrichment was quantified by comparing the frequency of motif occurrences in differentially accessible promoters relative to a background set consisting of all annotated promoters detected by ATAC-seq.

Statistical significance of motif enrichment or depletion was evaluated using Fisher’s exact test, comparing the number of promoters containing at least one motif occurrence in the differential set versus the background set. Resulting p-values were corrected for multiple hypothesis testing using the Benjamini–Hochberg procedure to control the false discovery rate (FDR). Motifs with FDR-adjusted p-values < 0.05 were considered significantly enriched or depleted. Enrichment effect sizes were expressed as odds ratios (OR), where OR > 1 indicates motif enrichment and OR < 1 indicates motif depletion.

Motif enrichment results were visualized using volcano plots displaying log odds ratio versus −log10(FDR), highlighting motifs associated with accessibility gains or losses.

### Chromosome-level density analysis of differential chromatin accessibility

To determine whether differential chromatin accessibility was preferentially enriched on specific chromosomes independently of baseline accessibility, chromosome-level density analysis was performed using the full ATAC-seq peak set as genomic reference. For each autosome, the total number of ATAC-seq peaks and the number of UP and DOWN DARs were quantified. Expected DAR counts were calculated based on the genome-wide proportion of differential accessibility relative to the total peak set, thereby controlling for chromosome-specific differences in peak density, chromosome length, and baseline accessibility. Enrichment or depletion was quantified by comparing observed and expected DAR counts and expressing the results as log₂-transformed observed-to-expected ratios. Statistical significance was assessed using hypergeometric tests, and resulting p-values were corrected for multiple testing using the Benjamini-Hochberg procedure to control the false discovery rate. Chromosome X was excluded from analysis due to sex-specific chromatin organization and mixed-sex cohort composition.

### Genome-wide hotspot analysis of chromatin accessibility remodeling

To identify localized genomic domains exhibiting clustering of chromatin accessibility changes, genome-wide hotspot analysis was performed using a window-based enrichment approach. The genome was partitioned into non-overlapping windows of 5 megabases using the T2T-CHM13 reference genome assembly. Within each window, the total number of ATAC-seq peaks and the number of UP and DOWN DARs were quantified. Expected DAR counts were calculated based on genome-wide DAR proportions, thereby normalizing for local peak density. Statistical enrichment of accessibility gains or losses was evaluated using hypergeometric testing, followed by Benjamini-Hochberg correction for multiple testing. Genomic windows with FDR-adjusted p-values below the significance threshold were defined as chromatin accessibility remodeling hotspots. This analysis enabled identification of spatially constrained genomic regions exhibiting coordinated accessibility remodeling.

### Nucleolus-associated domain enrichment analysis

To assess the relationship between chromatin accessibility remodeling and nucleolus-associated chromatin, DARs were intersected with nucleolus-associated domain (NAD) coordinates obtained from published NAD-seq datasets generated in HeLa cells (Peng et al. 2023). These datasets provide genome-wide maps of chromatin domains associated with the nucleolar periphery and enriched in heterochromatin and repeat elements. NAD coordinates were converted to the T2T-CHM13 reference genome assembly using LiftOver, and only successfully mapped intervals were retained. Overlap between NADs and ATAC-seq peaks was determined using GenomicRanges. Enrichment was quantified by comparing the proportion of DARs overlapping NAD regions to the corresponding proportion of background ATAC-seq peaks. Statistical significance was assessed using Fisher’s exact test, and p-values were corrected for multiple testing using the Benjamini-Hochberg procedure. Enrichment effect sizes were expressed as odds ratios.

To evaluate chromosome-specific architectural remodeling, enrichment analysis was additionally performed independently for each chromosome. Observed and expected overlaps between DARs and NAD regions were calculated relative to chromosome-specific peak distributions, and enrichment was quantified using log₂-transformed odds ratios. Statistical significance was assessed using Fisher’s exact test with correction for multiple testing across chromosomes.

### Lamina-associated domain enrichment analysis

To determine whether chromatin accessibility remodeling preferentially affects lamina-associated chromatin, DARs were intersected with lamina-associated domain (LAD) coordinates obtained from genome-wide DamID datasets generated in human fibroblasts (NKI fibroblast dataset, ENCODE)(Guelen et al. 2008). LAD coordinates were converted to the T2T-CHM13 genome assembly using LiftOver. Overlap analysis was performed using GenomicRanges, and enrichment was quantified by comparing DAR overlap frequency to the background ATAC peak set. Statistical significance was assessed using Fisher’s exact test followed by Benjamini-Hochberg correction. Chromosome-specific enrichment analysis was performed using the same approach, enabling identification of chromosomes exhibiting preferential association of chromatin accessibility changes with lamina-associated chromatin domains.

### Chromatin state enrichment analysis using ChromHMM

To determine the chromatin state context of accessibility remodeling, DARs were intersected with chromatin state annotations generated using ChromHMM (Ernst and Kellis 2012). ChromHMM segmentation datasets derived from ENCODE histone modification profiles were used as genome-wide reference maps of regulatory and repressive chromatin states. Because ChromHMM annotations are most consistently available in hg19 coordinates, genomic coordinates of ATAC-seq peaks and DARs were converted to hg19 using LiftOver prior to analysis.

Overlap between DARs and ChromHMM chromatin states was determined using GenomicRanges. Enrichment of DARs within individual chromatin states was quantified by comparing the frequency of state overlap in DARs relative to the background ATAC peak set. Statistical significance was assessed using Fisher’s exact test followed by Benjamini-Hochberg correction for multiple testing. Enrichment effect sizes were expressed as odds ratios. Chromosome-specific enrichment analysis was additionally performed to evaluate whether chromatin state associations were uniformly distributed or preferentially localized to specific chromosomes.

### Repeat annotation and enrichment analysis

To assess the association between chromatin accessibility remodeling and repetitive genomic elements, repeat annotations were obtained from the RepeatMasker track (https://www.repeatmasker.org/) corresponding to the T2T-CHM13 reference genome assembly. RepeatMasker annotations provide genome-wide classification of repetitive elements, including SINEs, LINEs, long terminal repeats (LTRs), DNA transposons, simple repeats, and low-complexity regions.

RepeatMasker annotation files were downloaded from the official T2T consortium genome annotation resources and imported into R as genomic intervals. Overlap between repeat elements and ATAC-seq peaks was determined using GenomicRanges.

Enrichment analysis was performed independently for UP DARs, DOWN DARs, and the full ATAC peak set used as genomic background. Enrichment was quantified by comparing the proportion of DARs overlapping each repeat class relative to background peaks.

Statistical significance was assessed using Fisher’s exact test followed by Benjamini-Hochberg correction to control the false discovery rate. Effect sizes were expressed as odds ratios. This analysis enabled identification of repeat classes preferentially associated with chromatin accessibility gains or losses.

### Analysis of canonical DUX4 target loci

A curated set of canonical DUX4-responsive genes was generated by integrating gene lists reported in multiple independent DUX4 induction and FSHD muscle studies (Geng et al. 2012; Young et al. 2013; Hendrickson et al. 2017; Jiang et al. 2020; Bosnakovski et al. 2023).

Gene symbols were harmonized using HGNC-approved nomenclature, and duplicate entries, transcript isoforms, and paralog redundancies were collapsed at the gene level. The resulting non-redundant list comprised 73 DUX4-responsive loci.

Because DUX4 target annotation varies across studies due to repeat-associated paralogs and subtelomeric gene clusters, this curated list was defined independently of expression in the present dataset to avoid circularity.

For each gene, genomic intervals spanning −2 kb to +2 kb relative to annotated transcription start sites (TSS) were defined using the T2T-CHM13 genome annotation. Overlap between these DUX4 target regions and ATAC-seq peaks or differentially accessible regions (DARs) was determined using the GenomicRanges package in R

Enrichment of accessibility gains or losses within the DUX4 target set was assessed relative to genome-wide promoter distributions using Fisher’s exact test, followed by Benjamini–Hochberg correction.

## Software availability

All analyses were performed using R (version 4.5.2) within the Bioconductor framework (Huber et al. 2015). Genomic interval operations were performed using the GenomicRanges package (Lawrence et al. 2013) and differential accessibility analyses were conducted using the csaw and edgeR packages (Lun and Smyth 2016; Robinson et al. 2010). Custom scripts used for genomic interval processing, enrichment analysis, and figure generation are available upon reasonable request.

## Data Access

All raw and processed sequencing data generated in this study have been deposited in the NCBI Gene Expression Omnibus (GEO) under accession number GSE322947. The data are available to reviewers using the token: ydanockcrvedzij.

## Acknowledgments

We are indebted to all patients and their families for their participation in this study. We thank Professor Paul D. Kaufman for helpful discussions and insightful comments on this work.

## Author contributions

F.L. and VS designed the research. F.L. and B.F. performed the research. F.L., V.S. and R.T. interpreted the results. VS and RT wrote the manuscript.

